# Growth response and recovery of *Corynebacterium glutamicum* colonies on single-cell level upon defined pH stress pulses

**DOI:** 10.1101/2021.02.26.433016

**Authors:** Sarah Täuber, Luisa Blöbaum, Volker F. Wendisch, Alexander Grünberger

## Abstract

Bacteria respond to pH changes in their environment via pH homeostasis to keep the intracellular pH as constant as possible within a small range. A change of the intracellular pH value influences e.g., the enzyme activity, protein stability, solubility of trace elements and the proton motive force. Here, the species *Corynebacterium glutamicum* has been chosen as a neutralophilic and moderately alkali-tolerant bacterium capable of maintaining an internal pH of 7.5 ± 0.5 in environments with an external pH between 5.5 and 9. In the recent years, the phenotypic response of *C. glutamicum* to pH changes has been systematically investigated at the bulk population level. A detailed understanding of the *C. glutamicum* cell responding to defined short-term pH perturbations/ pulses is missing. In this study, dynamic microfluidic single-cell cultivation (dMSCC) was applied to analyse the physiological growth response of *C. glutamicum* upon precise pH stress pulses at a single-cell level. Analysis of the growth behaviour at the colony level by dMSCC exposed to single pH stress pulses (pH = 4, 5, 10, 11) revealed a decrease in the viability with increasing stress duration. Colony regrowth was possible after increasing lag phases when stress durations were increased from 5 min to 9 h for all tested pH values. Furthermore, the single-cell analysis revealed heterogeneous regrowth of cells after pH stress, which can be distinguished into two distinct behaviours firstly, cells continue to grow without interruption after the pH stress, and secondly, some cells rest for several hours after the pH stress before they start to grow again after this lag phase. This study provides the first insights into the single-cell response to acidic and alkaline pH stress adaptation of *C. glutamicum*.

## Introduction

Various environmental fluctuations influence the growth and the physiology of bacteria (1). One decisive parameter is the pH value which has an impact on the solubility of nutrients and the cellular metabolism (2)–(4). The external pH fluctuation has an important effect of the intracellular pH value of the bacteria. A change of the intracellular pH value affects e.g. the enzyme activity, protein stability, solubility of trace elements and the proton motive force (PMF) (2)–(5). The regulation of the intracellular pH value is maintained by pH homeostasis (6). *Corynebacterium glutamicum* is a neutralophilic organism that can maintain the internal pH of 7.5 ± 0.5 in environmental pH fluctuations ranging between 5.5 and 9.0 (2). Outside this range the internal pH collapses and finally the pH homeostasis fails. In acidic environments, a significant amount of reactive oxygen species (ROS) is produced, which leads to oxidation of methionine and cysteine residues of proteins or iron sulfur clusters as well as to DNA damages (2). As a result, the metabolism may change, e.g., the iron starvation response is activated, consequently affecting the TCA cycle and NAD and methionine synthesis. Due to the reduction of the methionine synthesis, cysteine accumulates, which is toxic in acidic milieus (2). Another important mechanism for the pH homeostasis in acidic environments is the potassium uptake via potassium channels. This stabilises the PMF, which is essential for *C. glutamicum* growth in acidic and alkaline environments. This electrochemical proton gradient through the bacterial cell membrane is kept constant by ion transporters. The ATP synthase is driven by the PMF (3). In acidic environments, the gradient increases so that the electrochemical potential is adjusted by potassium flux (7). The associated genetic response involves RNA polymerase sigma factors, which regulates large genetic networks. The removal of sigma factors decreases growth and survival under any stress condition (8). In addition to inorganically acidic environments, pH shifts can also be induced by organic acids, which affect not only the H^+^concentration but also the available carbon source (8). In alkaline environments, much less is known for the molecular adaption mechanism of *C. glutamicum*. A MdfA homologue is missing and the involvement of further sodium proton antiporters in the pH reaction is unknown. Genes coding for homologous proteins of the amino acid decarboxylase, e.g. AdiCA, GadABC and CadAB, are also absent (2),(9). The generation of a considerable electrochemical potential across the cell membrane is critical for the entry of protons, typically via cation/ proton antiporter (3). Until now, Mrp-Typ Na^+^(Li^+^)/ H^+^antiporter 1 and 2 were found to contribute to resistance in alkaline environments (10). It was also postulated that the concentrations of proteins of the succinate dehydrogenase complex and F_0_F_1_-ATP synthesis are increased (11). The reader is referred to Guo et al. (12) for a detailed summary of the known mechanism and suggested strategies for *C. glutamicum* to cope with pH stress.

This knowledge is based on bulk population studies with microbial cells cultivated in small bioreactors or shaking flasks to investigate pH homeostasis (2),(8). This approach is blind to population heterogeneity since the behaviour of individual cells may be hidden behind the average value. Therefore, no representative information can be found for individual cells (13), and cell-to-cell heterogeneity remains not understood (14). Furthermore, traditional cultivation lacks the temporal precision and spatial resolution, e.g., due to slow mixing in larger volumes, to perform stress response experiments such as defined stress pulses or even the investigation of oscillating stress conditions (1).

Novel microfluidic tools allow to analyse cellular behaviour at the single-cell level and enable the cultivation of cells under precise environmental conditions (15),(16). Recently, a dMSCC setup was described that enables the cultivation of cells at dynamic environmental conditions (17). The dMSCC setup was used to analyse different oscillation frequencies that show the influence of various nutrient limitations on *C. glutamicum*. This dMSCC system was used here because it supports fast and precise medium changes and its high degree of parallelization provide sufficient data for rigorous statistical evaluation.

In this work, the influence of different pH values and pH pulses on the growth of *C. glutamicum* was systematically investigated. The dMSCC system allowed to investigate the behaviour of populations at the single-cell level as provided single-cell data for responses to pH stress pulses, which had not been analysed before. Besides continuous cultivation with pH values between 5 and 10, the responses to single stress pulse (4, 5, 10 and 11) with stress durations between 5 min and 9 h were investigated to determine how long single cells survive under these pH stress conditions. Furthermore, it was addressed how long it takes for individual cells to recover from different stress durations and it was determined, if all cells regrow homogeneously after pH stress with the same lag phase or whether different subpopulations can be identified, which differ in their adaptive response to the pH.

## Material and Method

### Pre-cultivation, bacterial strain, medium

In this study, the bacterial strain *C. glutamicum* WT ATCC 13032 was cultivated in CGXII at 30°C(18). Each medium component was autoclaved. Afterwards, the pH value of the medium was adjusted. Prior dMSCC medium was sterile filtered. For all microfluidic experiments CGXII without MOPS was used.

The overnight pre-cultures of *C. glutamicum* were inoculated from glycerol stock in 10 mL CGXII medium (with MOPS) in 100 mL baffled flasks on a rotary shaker at 120 rpm. Cells from the overnight culture were transferred to inoculate the second culture with a staring OD_600_of 0.05. When the culture reached an OD_600_of around 0.2 the cells were seeded in the microfluidic device. After seeding, the cells were dynamically perfused with CGXII medium at pH 7 and CGXII medium with varying pH value according to the protocol described further down.

### Chip preparation

For the PDMS soft lithography mould, a silicon wafer was fabricated. The detailed process steps for the wafer fabrication is shown in Täuber et al. (17).

The wafer was covered with PDMS in a ratio 10:1 (Sylgard 184 Silicone Elastomer, Dow Corning Corporation, USA). Afterwards, the wafer was degassed in an exicator for 20 minutes and backed for 2 hours at 80°C (Universal cupboard, Memmert GmbH, Germany). After this, the PDMS chip was cut out from the wafer, the in- and outlets were punched with a 0.75 mm biopsy puncher (Reusable Biopsy Punch, 0.75mm, WPI, USA) and cleaned with isopropanol three times. The cover glass (D 263 T eco, 39.5×34.5×0.175 mm, Schott, Germany) was cleaned just as the PDMS chip. The PDMS chip and the cover glass were O_2_ plasma (Femto Plasma Cleaner, Diener Electronics, Ebhausen, Germany) oxygenised for 24 seconds and were assembled. Afterwards, a 2-minute post bake at 80°C was performed to strengthen the bonding.

### Chip design

The chip design published in Täuber et al. (17) was used with two inlets for the different pH values (Fig. 1A). In between there are several arrays of monolayer cultivation chambers (80 µm × 90 µm × 750 nm) (Fig. 1B), whereby the different oscillation zones are separated from each other by a channel with a width of 400 µm. The supply channels have a height of around 10 µm and a width of 100 µm.

**Figure 1:**
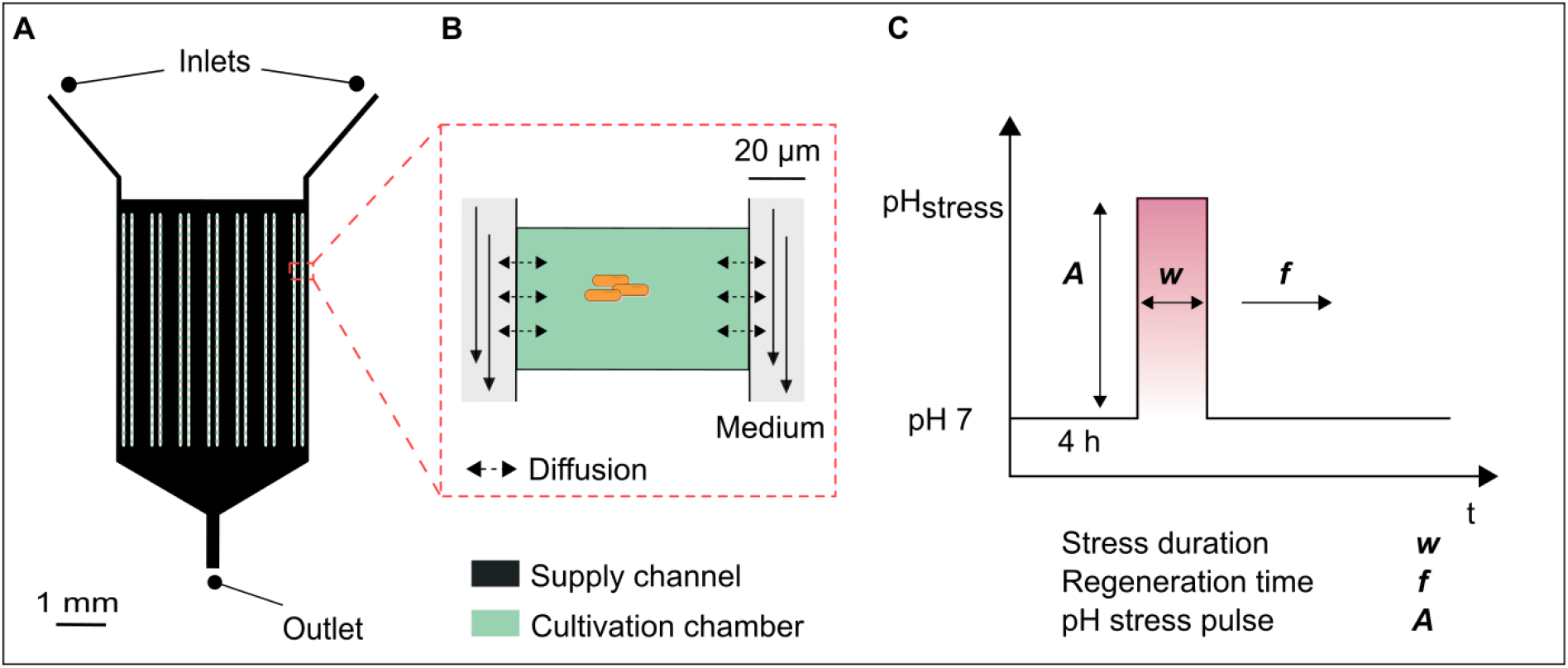
Design of the microfluidic device for the cultivation of single cells and small cell clusters under dynamic environmental conditions. A) dMSCC for single stress pulses. B) Monolayer cultivation chamber. C) Overview of parameters of dynamically controlled pH value.

### Setup and microfluidic cultivation

The culture preparation was described in “Pre-cultivation, bacterial strain, medium”. For the loading process, the cell suspension from the second culture was inoculated at an OD_600_of 0.2 in the microfluidic device. In a randomized process, about 75% of the cultivation chambers were loaded with ∼1-4 cells (19). When a sufficient number of chambers was filled with single cells, the flow of the cell suspension was stopped, and the medium flow was started. The medium flow was executed with high precision pressure pumps (Line-up series, Fluigent, Jena, Germany) with a pressure for the two-inlet chip of 180 mbar and 20 mbar to start the single stress pulses experiments (detailed flow profiles see Fig. S1). The pulse profiles were applied using a tailor made cultivation profile implemented into an automated software tool (microfluidic automation tool (MAT), Fluigent, Jena, Germany).

In this study, the cells were first adapted for 4 h in the microfluidic device at pH 7 before the pH stress was performed. The pH stress pulses (*A*) were varied between 4, 5, 10 and 11 for different experiments. Single stress pulses were performed with varying stress durations (*w)* between 5 min and 9 h (Fig. 1C). Afterwards, a regeneration time (*f*) was performed until the end of cultivation to analyse the regrowth of cells. During the measurements of the alkaline stress pulses, crystallization sometimes occurred at the boundary layer between the reference medium pH 7 and the alkaline pH values 10 and 11. Magnesium sulphate or magnesium hydroxide probably crystallized out at this boundary layer. The solubility of magnesium hydroxide is 9 mg/L, while the CGXII medium contains about 10 times this amount. The data shown here were free of crystals. The analysis of the experiments was stopped, when crystallization occurred and the flow profile was significantly altered, to guarantee a proper interpretation of the data.

### Live-cell imaging

Live-cell microscopy was conducted using an inverted automated microscope (Nikon Eclipse Ti2, Nikon, Germany). The microscope was equipped in a cage incubator for optimal temperature control (Cage incubator, OKO Touch, Okolab S.R.L., Italy). The microfluidic chip was attached to an in-house manufactured holder. The setup was equipped with a 100x oil objective (CFI P-Apo DM Lambda 100x Oil, Nikon GmbH, Germany), a DS-Qi2 camera, and an automatic focus system (Nikon PFS, Nikon GmbH, Germany) to prevent the thermal drift during cultivation. 100 cultivation chambers were manually selected in each experiment using the NIS Elements software (Nikon NIS Elements AR software package, Nikon GmbH, Germany). The images were taken every 10 minutes with an exposure time of 50 ms.

### Image and data analysis

Image analysis of the image sequences was done using the open source software Fiji (20). In the phase contrast images the cells were separated from the background for each time point with the k-mean clustering for background correction. The cluster with the cells was maintained and the background and the intermediate area between the cells were deleted. Cells that were located too close together could not be separated by clustering. Therefore, a watershed transformation was performed. The integrated analyse particle was then used to determine the cell count and cell area for each time point.

Using OriginPro 2019b (OriginLab Corporation, Northampton, USA), the growth curves were plotted, and the colony growth rate was determined by a linear fit through the semi-logarithmic plot of the cell number. In the case of no exponential growth, the colony growth was calculated over an increase of the cell area. Dynamic single-cell data illustrated in Figure 4 and 5 were analysed manually.

## Results

### C. glutamicum growth at constant pH environments

In a microscopy setup, the dMSCC approach allows to monitor single cells growing to colonies of up to 1000 cells in cultivation chambers with supply of cultivation medium. Colony growth is here defined by the increase in cell number or cell area in a cultivation chamber that typically started with a one or a few single cells at the beginning of cultivation. In the first set of experiments, microbial colony growth was investigated by dMSCC at different, but constant pH values ranging from 5 to 10. Colony growth was monitored by live-cell imaging (Fig. 2A) and pH specific growth rates for each pH were determined by cell number for growth rates above 0.15 h^-1^and by cell area for growth rates below 0.15 h^-1^(Fig. 2B). Two set of experiments were performed, the first one using HCl & KOH for pH adjustment and the second using NaOH & H_3_PO_4_. Based on the growth rates, a pH optimum curve for *C. glutamicum* on CGXII was derived with the influence of the acid or base used for pH adjustment (Fig. 2C).

**Figure 2:**
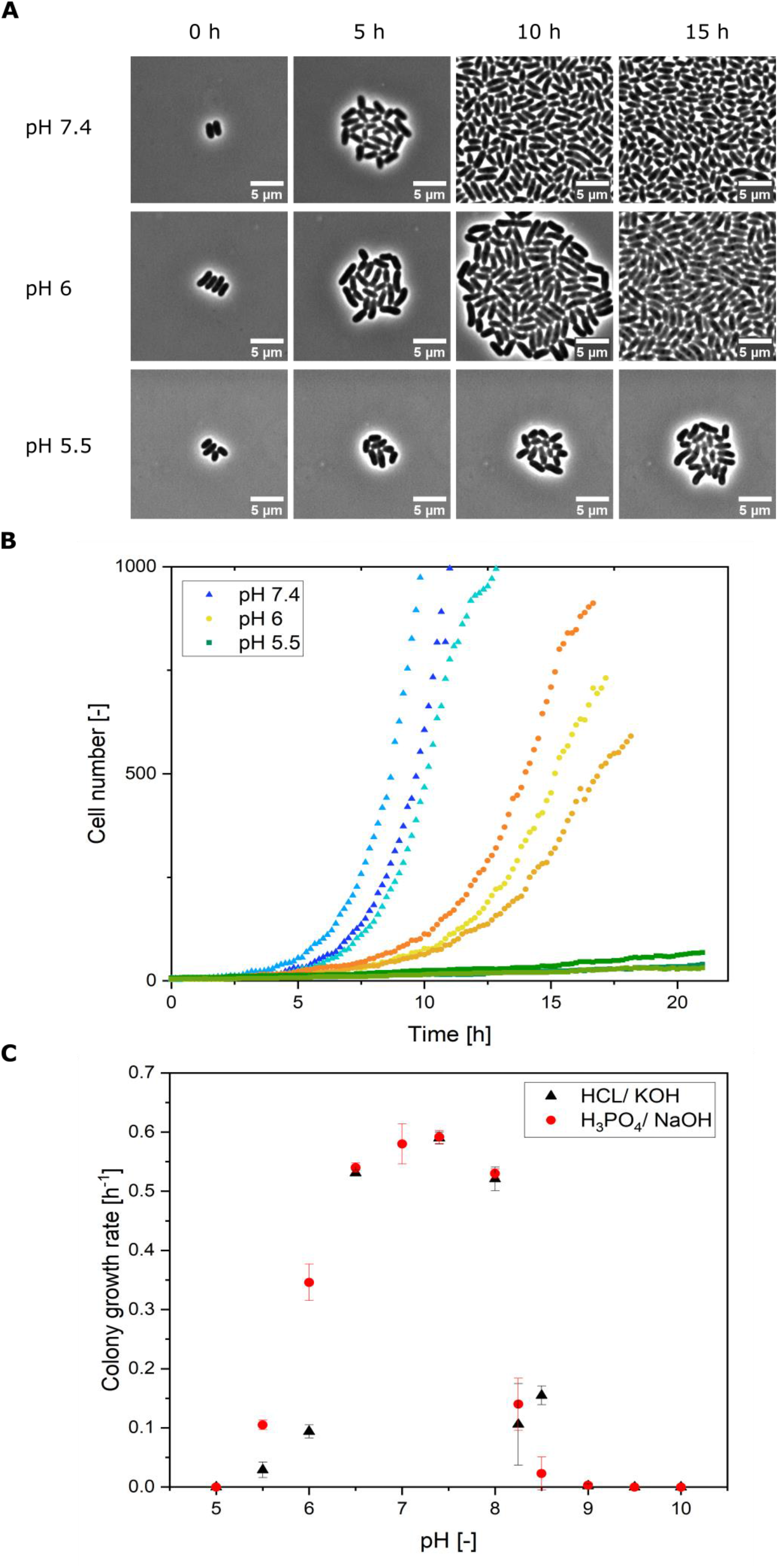
A) Microscope images of growing microcolonies in continuous pH conditions of pH 7.4, 6 and 5.5 at different points in the cultivation time. Scale was set to 5 μm. B) Colony growth of *C. glutamicum* at different, constant pH values (5.5, 6, 7.4) using NaOH & H_3_PO_4_ for pH adjustment. Three microcolonies are shown for each pH value. C) pH optimum curve for the colony growth of *C. glutamicum* in microfluidic chip on CGXII medium.

Optimal growth rates were obtained between pH 7 and 7.4 with a maximum growth rate of µ_max_= 0.59 ± 0.01 h^-1^ at pH 7.4. Exponential growth was observed for all microcolonies at pH values between pH 6 and 8. Microcolonies cultivated at pH 5 < 6 and 8 < 9 grew very slowly, but exponential growth was not observed. A linear growth of the cell area was monitored. Consequently, under these conditions the growth rates could not be determined quantitatively and were indicated as ≤ 0.1 h^-1^. At pH values ≤ 5 and ≥ 9 no growth was monitored. The choice of the acid or based used to control the medium pH did not influence the colony growth rate in a range of pH 6.5 to 8. However, at the very acidic pH values 5.5 and 6, growth rates were reduced by 27% when the pH of CGXII medium was adjusted with hydrochloric acid compared to phosphoric acid. A similar effect was observed at the alkaline pH value of 8.5, where cells in NaOH adjusted medium grew 82% slower than cells in KOH adjusted medium. H_3_PO_4_and NaOH were chosen for the next experiments, as both are also used in the bioreactor (21).

### C. glutamicum growth at single pH stress pulse experiments

After having studied growth of *C. glutamicum* at constant pH values, the response to a single pH stress pulse was studied. Therefore, pH stress amplitudes *A* between 4 and 11 and various stress durations *w* were tested when a single pH stress pulse was applied after 4 hours of growth in the dMSCC at pH 7. The aim was to answer the key question how *C. glutamicum* responds to abrupt pH changes of different amplitude and stress duration. Therefore, it was determined if cells continued to grow or stopped during the stress pulse, and if so, when regrowth after the pH stress pulse occurred (lag phase). The colony lag phase is defined here as the time between the end of the pH stress pulse and the onset of regrowth of a colony. Viability was defined as the ratio of cells showing regrowth and cell division after the stress pulses. Furthermore, the time until the first post-stress cell division (first division after the stress pulse means individual lag phase) was determined.

### C. glutamicum growth at the colony level after various pH stress pulses

In a first set of experiments colony growth upon pH stress pulses between 5 min and 9 h was determined. During the pH stress pulse, the colony area neither decreased nor increased (Fig. S2). After the pH stress pulse, an increase of the colony area was observed. Lag phase and viability after pH stress pulses with amplitudes ranging from 4 to 11 are shown in Figure 3. Colonies survive stress pulses of pH 5 and 10 with durations longer than 6 h (Fig. 3B&D). The lag phase after pH 5 stress pulses increased with increasing stress duration. Upon stress duration between 2 h and 4 h the lag phase was only two hours, after 9 h stress the lag phase was 12 h. The viability decreased drastically to 30% after 4 hours, after 9h stress the viability was only 6%. Thus, a relation between stress duration and elongation of the lag phase as well as loss of viability was found (Fig. S3). Upon pH 10 stress pulses, the lag phase increased from 2.8 to 18 h with increasing stress duration. Due to precipitation or crystallization at alkaline pHs, the values for pH 10 stress durations of 8 and 9 h could not be measured (see materials and methods). The viability decreased with increasing stress duration for pH 5 and pH 10 stress pulses. Upon pH 4 and 11 stress pulses, the viability decreased faster. Already after 30 min no viable cells were found (Fig. 3A&C). At pH 4, after 5 min of stress the cells continued to grow without lag phase and the viability was close to 100%. After 10 min stress pulse, the viability decreased to only 20% while at pH 11 the viability decreased to only ∼70%. Fig. 3 shows two main results concerning the viability of the colonies: First, the viability decreased between 2 and 6 h after stress duration with pH 10 (Fig. 3D), at longer stress durations viability remained constant at around 20%. Second, a rapid decrease of the viability can be seen at pH 4 between 0 and 5 min to 20% (Fig. 3A). Afterwards, the viability decreased continuously to 0% after 30 min of stress.

**Figure 3:**
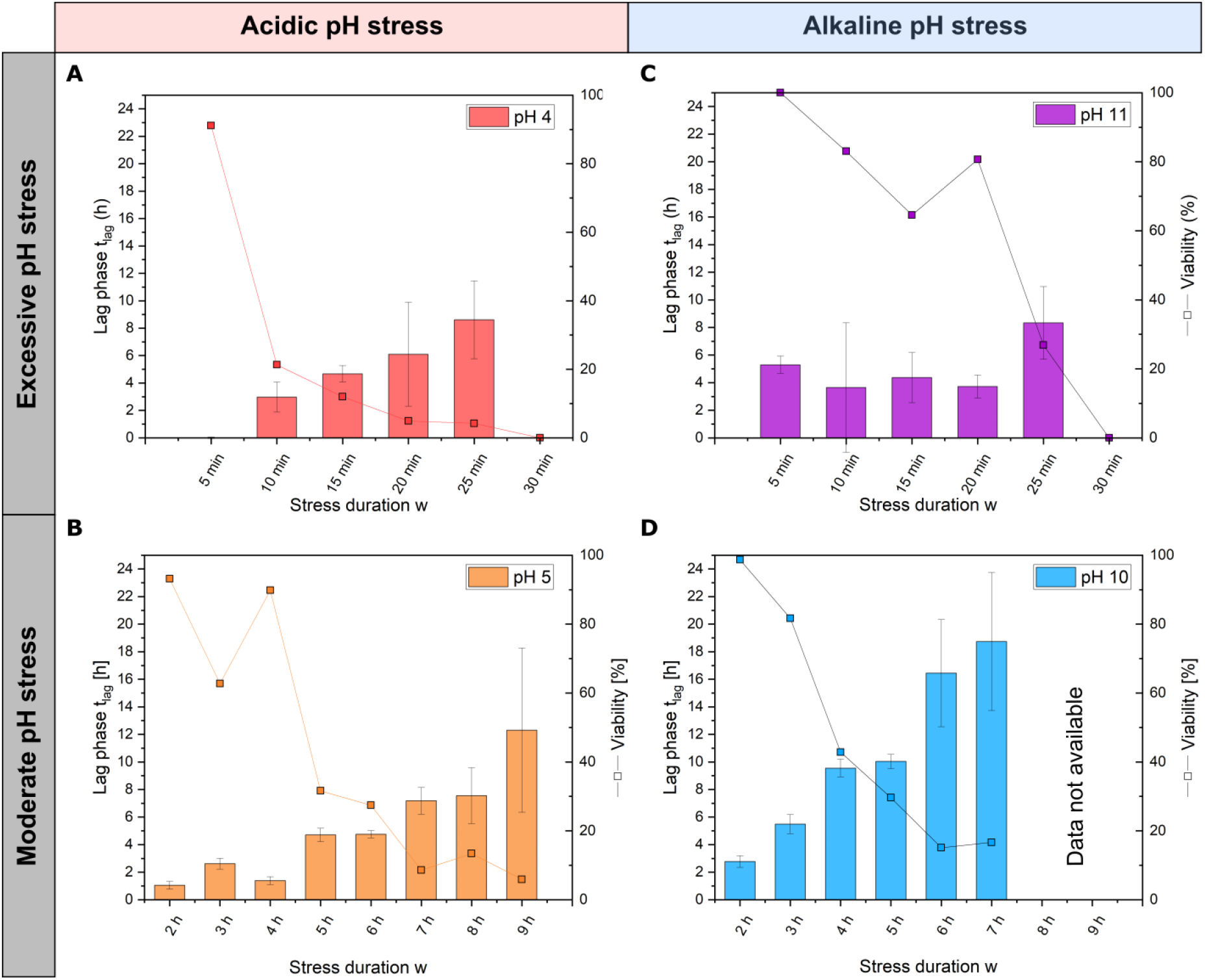
Lag phase and viability of *C. glutamicum* at different pH stress pulses. Lag phase and viability at acidic A) pH 4 and B) pH 5 stress pulses and at alkaline C) pH 11 and D) pH 10 stress pulses.

### C. glutamicum growth at the single-cell level after various pH stress pulses

To gain insight whether *C. glutamicum* shows a homogenous response to pH stress pulses or whether heterogeneity and distinct sub-populations arise, the single-cell behaviour upon pH stress pulses was analysed by measuring the time until the first post-stress division event (= individual lag phase) after the different stress durations *w* and different pH stress amplitude *A*.

With acidic pH values, the first post-stress division times show significant variation (Fig. 4A&B). At pH 5 some cells divided directly after the stress phase (stress durations between 2 and 6 h) while other cells in the same colony only divided after a lag phase. After 2 h stress duration, a normal distribution of the first post-stress division was observed as expected for a homogeneous population (see Fig. S4A). The longer the duration of the stress, the larger the median of the first post-stress division and the broader was their distribution. After 6 h stress duration, no normal distribution was observed, the distribution was wider, indicating two subpopulations (see Fig. S4B). At pH 4, the first post-stress division increased with increasing stress duration so that at low stress durations single cells divided in less than 2 h. After 25 min, only a few cells survived (Table S1), and for the first post-stress division a large variation between 1 h and 17 h was observed.

**Figure 4:**
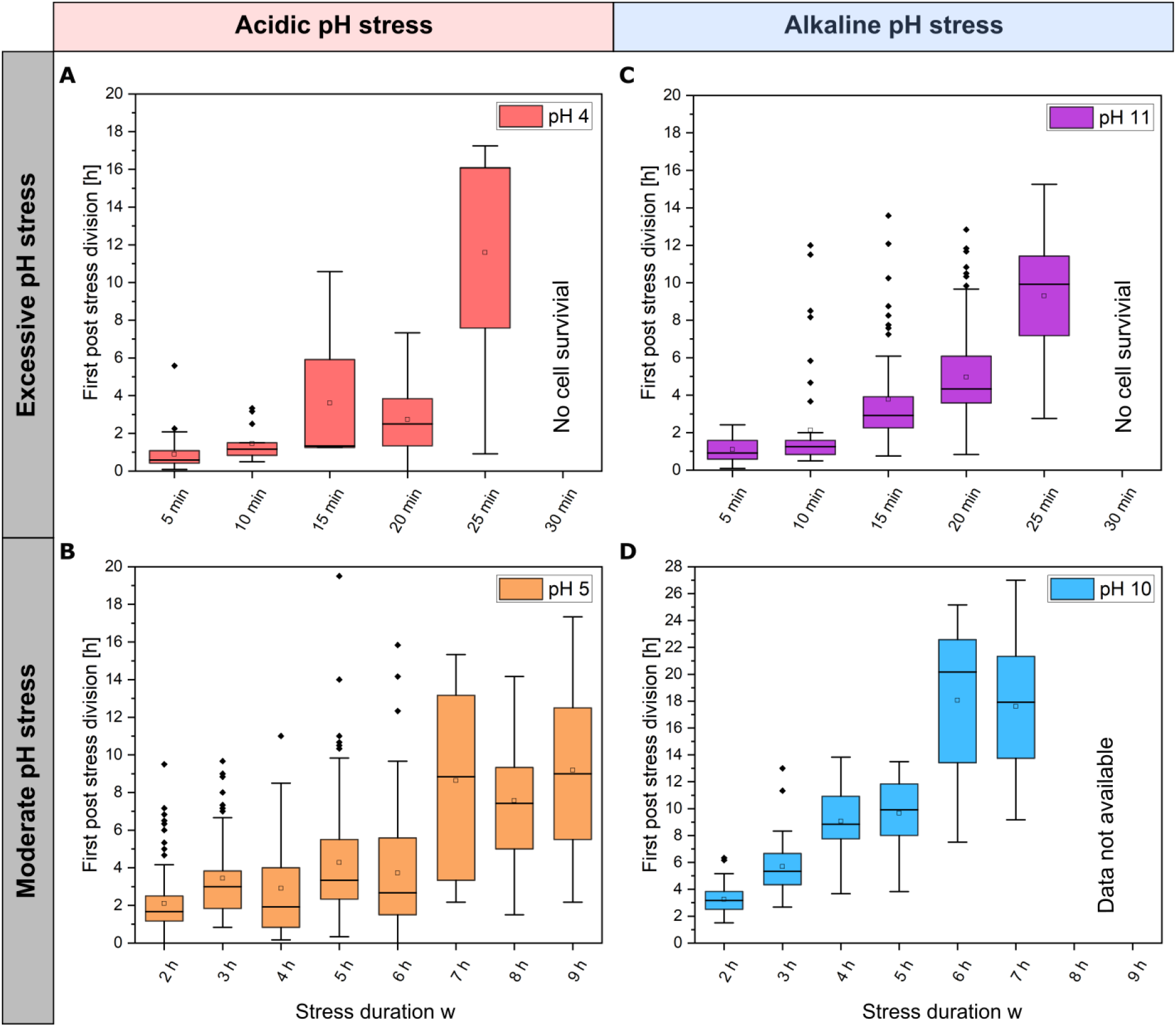
Distribution of the first post-stress division of a single cell after a pH stress pulse of A) pH 4. B) pH 5. C) pH 11. D) pH 10. Overview of the sample size and the surviving single cell numbers is shown in Table S1.

In alkaline stress pulses at pH 11, a great variation of single cell post-stress division was observed which ranged from 1 h up to 12 h. After 30 min of pH 11 no cells survived (Fig. 4C). At pH 10, stress pulses revealed that the first division after a stress pulse required between 3 h and 27 h positively correlating with stress duration (Fig. 4D).

### Single-cell growth behaviour and adaptation during the first post-stress division

After investigating the time up to the first post stress division, the regrowth behaviour of single cells until the first post-stress division was studied in more detail, i.e., including cell elongation prior to division. Motivating questions were: Are there different adaptation phenotypes upon pH stress? Do cells stay dormant and restart growth with pre-stress growth rates, or do cells restart with a slower growth rate?

The elongation behaviour of individual cells was examined by measuring the cell length over time until the first post-stress division after a single pH stress pulse. Four different phenotypes were found at pH 5 (Fig. 5A&C): first, cells that started their exponential elongation immediately after a stress duration of 2 and 6 h (Fig. 5E&G – cell number A.2). Second, cells that showed a linear/ slow elongation with no lag phase (Fig. 5A&E&G – cell number A.3). Third, some cells showed a lag phase ranging in duration from minutes to several hours before cell elongation started (Fig. 5A&E&G – cell number A.1). Fourth, cells that failed to elongate after 2 h of stress pulse of pH 5 (see Tab. S1 – cells not shown in Fig. 5). After a stress duration of 9 h (Fig 5E), three phenotypes (Phenotype 2-4) were observed. The cells showed a broadly distributed lag phase. Three phenotypes were found at pH 10 (Fig. 5B&D): first, cells showing a lag phase before elongation (2 to 14 h lag time) (Fig. 5F). These cells remain dormant for up to 14 hours until they begin to elongate exponentially and divide approximately 19 h after pH stress (Fig. 5D&F&H – cell number B.2 and Fig. S5). Other cells started their elongation after a short growth lag phase (Fig. 5D&F – cell number B.1). The second phenotype shows cells that elongate linearly after a lag phase (Fig. 5F&H – cell number B.3). Regardless of the growth phenotype, except for the first post-stress division, all daughter cells were viable and have a division time of 69.7 ± 6.3 min (see Fig. S6). Third, 90 cells failed to survive the 6 h stress pulse at pH 10 (see Tab. S1 – cells not shown in Fig. 5).

**Figure 5:**
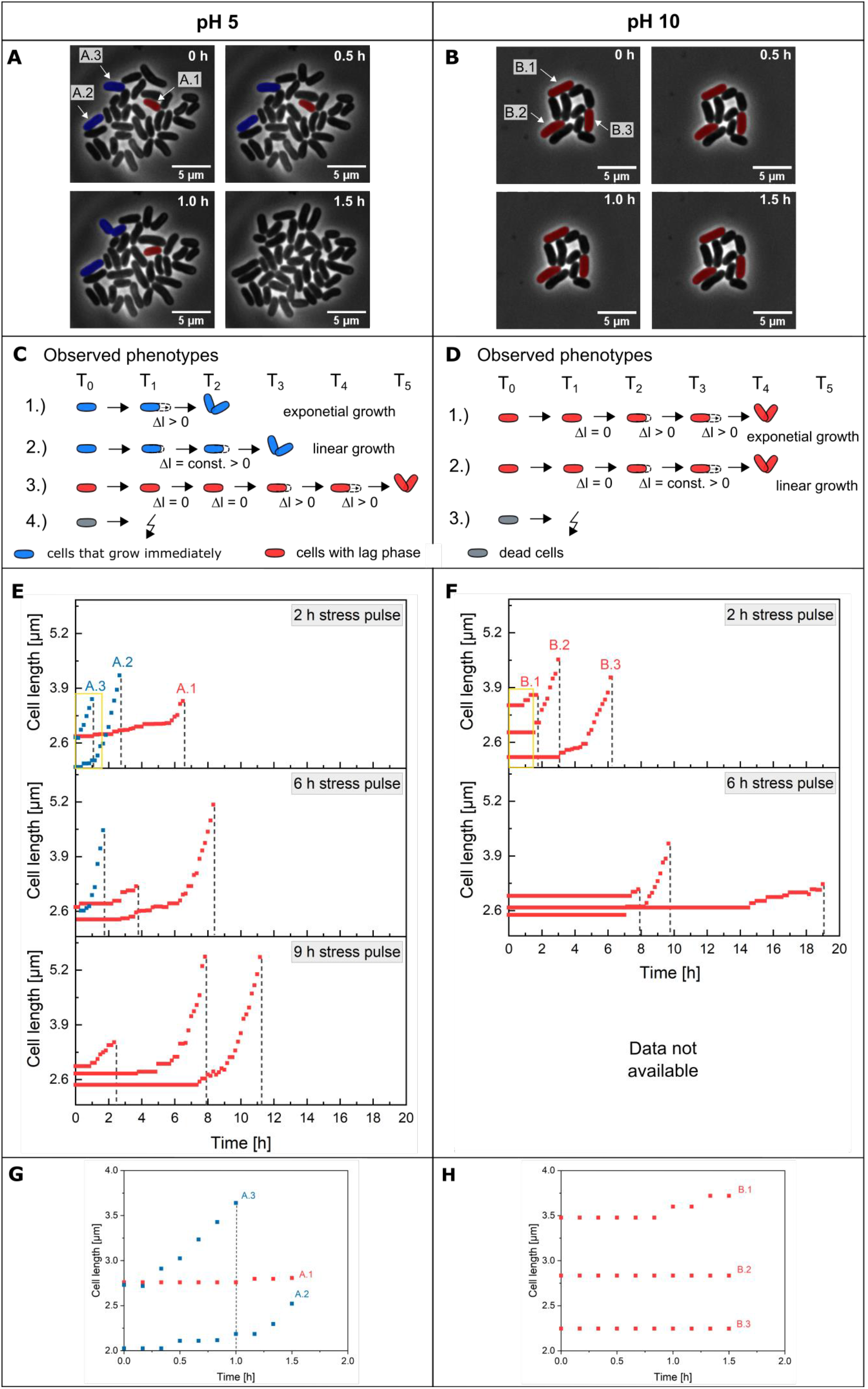
Single-cell adaptation of *C. glutamicum* after pH stress pulses. Microscope images of growing colonies after 2 h stress pulse of A) pH 5 and B) pH 10 at different post stress time periods. Cells marked blue could be preadapted cells and red marked cells are not adapted cells showing a lag phase before elongation and division. Scale was set to 5 μm. Observed phenotypes are shown in C) for pH 5 and D) for pH 10. E)&F) Cell elongation of single *C. glutamicum* cells after different stress pulses (2 h, 6 h and 9 h) of pH 5 and 10 stress until the first post-stress division. The dashed line marks the first post-stress division of the cells. G)&H) Zoom-out of the yellow box in C)&D) at the first 1.5 h after a 2 h stress pulse. Growth behaviour until the first post-division are shown.

## Discussion

### C. glutamicum growth at constant pH environments

The results obtained with dMSCC of *C. glutamicum* growth at constant pH confirm former experiments obtained in bulk scale (2),(8). Jakob et al. found a comparable limiting pH range from 5.5 to 9.5 in a turbidostat fermenter using lactic acid and NaOH for pH control. Follmann et al. reported a slightly broader pH range of 4 to 10. This difference may be due to batch cultivation condition in their study compared to the experiments performed under continuous conditions in this study. Batch cultivations are typically characterized by changing conditions as, e.g., the limiting substrate is consumed, products and by-products accumulate. It is possible that *C. glutamicum* exports ions and metabolites that promote growth at stress pH during a batch cultivation (12). These effects are minimal in the setup used here. Consequently, the maximum growth rate of *C. glutamicum* reported here exceeded that observed in shake flasks or bioreactors. The maximum growth rate of µ_max_ = 0.59 ± 0.01 h^-1^at pH values of 7 to 7.4 (Fig. 2C) is comparable to that measured with a similar setup, namely in a perfusion based microfluidic single-cell cultivation system (22).

The finding that pH adjustment with different bases had a significant influence on the growth rate, i.e., faster growth when the pH was adjusted with KOH and H_3_PO_4_may be explained by the fact that pH control alters ion concentrations in the medium, which may affect cells by different transmembrane ion gradients. With a concentration between 200 and 800 mM, K^+^is the most abundant ion in the cytoplasm of *C. glutamicum*. K^+^ions have been shown to be important for its acidic stress tolerance (7), whereas under alkaline conditions a higher K^+^concentration has no effect on the growth (7). The *C. glutamicum* mineral salts medium has a very low concentration of Na^+^ions because they have a negative effect on growth. The Na^+^/ H^+^antiporter Mrp1 maintains a low intracellular Na^+^ ion concentration. Deletion of its gene or chromosomal replacement of lysine 299 in the Mrp1A subunit increased the intracellular Na^+^ion concentration and led to a more alkaline intracellular pH value, which attenuated growth strongly (10).

### C. glutamicum growth at single pH stress pulse experiments

Detailed analyses of survival after pH stress pulses are missing in the literature for *C. glutamicum*, but *E. coli* and *B. subtilis* have been studied to some detail (23)–(25). To perform these analyses at the single cell level, appropriate tools have been missing so far. These few reported *E. coli* studies are difficult to compare with this study because they did not follow cell elongation and division but monitored how the cytoplasmic pH changed and recovered after shifting the extracellular pH. Moreover, the applied stresses involved only small pH shifts of 0.5 to 2.0 units. Martinez et al. (25) observed the cytoplasmic pH after an acidic pH shift using ratiometric pHluorin and they showed that pH homeostasis varied between the studied bacteria. An external pH shift from pH 7.5 to 6 revealed a recovery of the cytoplasmatic pH within 7 minutes after the shift for *E. coli* with only 3% failing to recover. The cytoplasmatic pH of *E. coli* decreased by 1.5 pH units as consequence of shifting the external pH from 7.5 to 5.5. The cytoplasmic pH started to recover after 10 seconds, however, this process was biphasic with a rapid recovery for 0.5 minute followed by a gradual recovery for 4 min until a pH of 7.4 was reached (24),(26). Wilks et al. (23) analysed the growth of *B. subtilis* after pH shifts and found that after a pH shift from 8.5 to 6 cells started to grow rapidly again with a short lag phase. However, after a pH shift from 6 to 8.5 *B. subtilis* showed a long lag phase. Similar to the results obtained with *B. subtilis* our study also found long lag phases of several hours after an alkaline stress that increase with the stress duration.

For all pH stress pulses, lag phases did not only increase with increasing stress pulse duration, but a broadening of the first-post stress division was also found, which may indicate increasing heterogeneity of coping with pH stress by genetically identical cells. At pH 4 and 11, a strong effect was observed after the pH stress due to the rapid decrease of the viability. After 30 min of pH 11 no cells survived. One possible mechanism could be mycothiol, which protects cysteine and methionine residues of proteins from ROS by reversible S-mycothiolations. This allows for fast protein regeneration after acidic stress (27). In alkaline environments no ROS protection is necessary (2).

The reduction of the viability of pH 4 and 11 could be similar to the thermal death curves observed for various bacterial species. Viability at pH 4 decreased at a similar rate as found in thermal death curves for *Cl. botoulinum*, for which a rapid decrease (lower than 2 minutes) in cell survival at 121°C was observed (28). For *B. sporothermoduranes* a slower decrease of the survival curve at 121°C was found, similar to the decrease of viability observed here at pH 11. For *C. glutamicum* no studies on thermal death points could be found. pH 4 and 11 could thus have a disinfecting effect within a culture.

In the last part of the study, cell elongation prior to the first post-stress division was investigated in more detail. Four different phenotypic subpopulations were found after a pH 5 stress pulse. Possibly, the preculture may already have contained these subpopulations that differ in their preadaptation to pH stress. Alternatively, the four subpopulations may have arisen stochastically. In the first subpopulation (see Fig. 5C-1&2), cells do not seem to be significantly affected by the stress pulse and started to elongate immediately after the stress pulse. These cells may have been preadapted to the environmental changes so that no damage occurred from the pH stress (Fig. 5C – blue cells). The cells of the second subpopulation might not have been adapted to the environmental changes and showed a lag phase. After the growth lag phase, the cells regain maximal growth rate indicating, that full cell integrity is obtained after a short adaption phase. In this case, proteins may be denatured, the PMF is compromised, and further damage from pH stress may occur, requiring time to repair. In the third subpopulation, the dying cells may have a compromised PMF and further damage due to pH stress that is unrepairable (Fig. 5C-4). These cells may also be dormant, so that extra addition of growth-enhancing substances can cause the cells to start growing again. *C. glutamicum* has two open reading frames, rpf1 and rpf2, which are similar to the essential resuscitation promoting factor (Rpf) from *M. luteus*, which would enable renewed growth through the addition of additional substances (29),(30). At pH 10, a phenotypic heterogeneity is evident because of the different lag phases. Similar phenotypes to the acidic stress were observed, with one difference: all phenotypes have a growth lag phase (Fig. 5D-1-3).

Especially under stress conditions, subpopulations with genotypic or phenotypic adaptation mechanism are beneficial for the survival of a population (31),(32). In alkaline pH, compared to acidic stress, there is a growth lag phase (no direct regrowth observed) in which the cells do not grow and stagnate for several hours before length growth was observed. Here, it is possible that a genetic response to the pH stress is necessary, which requires time. As for example in case of osmotic stress (33), first a fast response by transport and uptake of compatible substances from the medium is carried out and then a genetic response is triggered. Acidic stress may require only biochemical reaction mechanism (e.g., mycothiol, protein refolding, iron transport, etc.) to respond to the stress pulse.

The observed phenotypes need to be further examined in future studies, e.g., with a fluorescence signal that is coupled to PMF (34), so that loss of capability or damage during or after pH stress can be analysed in more detail. Furthermore, detection of the cytoplasmic pH can be determined with pH sensitive fluorescent proteins (6). Further insights into the heterogeneity and adaptive behaviour in response to pH stress pulses could be gained from fluorescence-based sensors that can report internal cellular pH values (35),(36). Non-invasive detection of the cytoplasmic pH could answer the questions about how long *C. glutamicum* is able to maintain the homeostasis and how physiology is affected upon pH stress pulses in the moment the pH homoeostasis collapses.

Further and detailed insights into heterogeneity within single-cell adaptation after pH stress pulses could be obtained by using single-cell growth channels, often referred to as mother machines (37), instead of the cultivation chamber used in this work. Here, single cells can be grown over several generations and detailed dynamic single-cell studies can be performed. This allows quantification of single-cell growth (i.e., elongation and time until cell division) over many generations and allows determination of whether and to what extend the pH stress pulse has an effect on, for example, maximum growth rates on successive generations following the stress pulse. In future, these systems could even be used to investigate cellular behaviour of single cells upon oscillating pH stress or upon multi (two or three pulses)-pH stresses (38). Here, questions such as how individual cells respond to oscillating pH stress can be answered.

## Conclusion

This study investigates the influence of different pH pulses on single-cell level. The growth response of *C. glutamicum* on single-cell level was analysed at different single stress pH values using dMSCC. This study confirms the results from bulk scale experiments at constant pHs. Long-term cultivations at non-optimal pH values were performed to analyse the growth range between 5 and 10 of *C. glutamicum*. Colony growth in the range of pH 6 to 8 was observed, which is consistent with lab-scale systems. New insights were shown on the growth behaviour during non-optimal pH and different pH stress pulses, which had not been possible before. Different single stress pulses of pH values between 4 and 11 were performed to analyse how long cells could survive in this pH values and how heterogeneously the cells continue to grow after the stress pulses. *C. glutamicum* can survive up to 9 h in pH 5, the variability decreases continuously after 5 h stress up to 6% after 9 h stress. At pH 10 the viability is constant at 20% after 6 h of stress. It has been shown that a growth lag phase after acidic stress is shorter than at alkaline stress. In addition, the growth behaviour of single cells after stress duration was analysed. It was found that the first post-stress division varies strongly at all pH values, indicating heterogeneous behaviour. Four phenotypes were observed at acidic pH values: first and second, cells are preadapted to the stress pulse to a stress duration of 6 h and length growth could be exponential or linear. Third, cells show a lag phase before the cell elongation and division begins. Fourth, cells do not survive the stress pulse. At alkaline pH values only three phenotypes were found: first and second, cells stagnate for several hours until (exponential or linear) elongation starts. Third, cells failed to survive the stress pulse. This study provided the basis for microbial phenotypes and adaptation upon pH stress pulses and lays the foundation for further studies under fluctuating and oscillating pH conditions.

## Acknowledgements

Parts of this work were performed at the cleanroom facilities of the Department of Biophysics and Nanoscience as well as the Department for Physics of Supramolecular Systems and Surfaces at Bielefeld University. The authors are thankful for all the help and support.

## References

1. Lara AR, Galindo E, Ramírez OT, Palomares LA. 2006. Living With Heterogeneities in Bioreactors: Understanding the Effects of Environmental Gradients on Cells. Molecular Biotechnology 34, 355–382. doi: 10.1385/MB:34:3:355.

2. Follmann M et al. 2009. Functional genomics of pH homeostasis in Corynebacterium glutamicum revealed novel links between pH response, oxidative stress, iron homeostasis and methionine synthesis. BMC Genomics 10, 621. doi: 10.1186/1471-2164-10-621.

3. Krulwich TA, Sachs G, Padan E. 2011. Molecular aspects of bacterial pH sensing and homeostasis. Nat Rev Microbiol 9, 330–343. doi: 10.1038/nrmicro2549.

4. Rosenthal K, Oehling V, Dusny C, Schmid A. 2017. Beyond the bulk: disclosing the life of single microbial cells. FEMS Microbiol Rev 41, 751–780. doi: 10.1093/femsre/fux044.

5. Olson ER. 1993. Influence of pH on bacterial gene expression. Mol Microbiol 8, 5–14. doi: 10.1111/j.1365-2958.1993.tb01198.x.

6. Haynes EP et al. 2019. Quantifying Acute Fuel and Respiration Dependent pH Homeostasis in Live Cells Using the mCherryTYG Mutant as a Fluorescence Lifetime Sensor. Anal Chem 91, 8466– 8475. doi: 10.1021/acs.analchem.9b01562.

7. Follmann M et al. 2009. Potassium transport in corynebacterium glutamicum is facilitated by the putative channel protein CglK, which is essential for pH homeostasis and growth at acidic pH. J Bacteriol 191, 2944–2952. doi: 10.1128/JB.00074-09.

8. Jakob K et al. 2007. Gene expression analysis of Corynebacterium glutamicum subjected to long-term lactic acid adaptation. J Bacteriol 189, 5582–5590. doi: 10.1128/JB.00082-07.

9. Kalinowski J et al. 2003. The complete Corynebacterium glutamicum ATCC 13032 genome sequence and its impact on the production of l-aspartate-derived amino acids and vitamins. J Biotechnol 104, 5–25. doi: 10.1016/s0168-1656(03)00154-8.

10. Xu N et al. 2018. The Lysine 299 Residue Endows the Multisubunit Mrp1 Antiporter with Dominant Roles in Na+ Resistance and pH Homeostasis in Corynebacterium glutamicum. Appl Environ Microbiol 84. doi: 10.1128/AEM.00110-18.

11. Barriuso-Iglesias M, Schluesener D, Barreiro C, Poetsch A, Martín JF. 2008. Response of the cytoplasmic and membrane proteome of Corynebacterium glutamicum ATCC 13032 to pH changes. BMC Microbiol 8, 225. doi: 10.1186/1471-2180-8-225.

12. Guo J et al. 2019. Recent advances of pH homeostasis mechanisms in Corynebacterium glutamicum. World J Microbiol Biotechnol 35, 192. doi: 10.1007/s11274-019-2770-2.

13. Lindström S, Andersson-Svahn H. 2010. Overview of single-cell analyses: microdevices and applications. Lab Chip 10, 3363–3372. doi: 10.1039/c0lc00150c.

14. Lindemann D, Westerwalbesloh C, Kohlheyer D, Grünberger A, Lieres E von. 2019. Microbial single-cell growth response at defined carbon limiting conditions. RSC Advances 9, 14040–14050. doi: 10.1039/c9ra02454a.

15. Grünberger A, Wiechert W, Kohlheyer D. 2014. Single-cell microfluidics: opportunity for bioprocess development. Curr Opin Biotechnol 29, 15–23. doi: 10.1016/j.copbio.2014.02.008.

16. Dusny C et al. 2015. Technical bias of microcultivation environments on single-cell physiology. Lab Chip 15, 1822–1834. doi: 10.1039/C4LC01270D.

17. Täuber S, Golze C, Ho P, Lieres E von, Grünberger A. dMSCC: A microfluidic platform for microbial single-cell cultivation under dynamic environmental medium conditions (2020).

18. Unthan S et al. 2014. Beyond growth rate 0.6: What drives Corynebacterium glutamicum to higher growth rates in defined medium. Biotechnol Bioeng 111, 359–371. doi: 10.1002/bit.25103.

19. Probst C et al. 2015. Rapid inoculation of single bacteria into parallel picoliter fermentation chambers. Anal. Methods 7, 91–98. doi: 10.1039/C4AY02257B.

20. Schindelin J et al. 2012. Fiji: an open-source platform for biological-image analysis. Nat Methods 9, 676–682. doi: 10.1038/nmeth.2019.

21. Seletzky JM et al. 2007. Scale-up from shake flasks to fermenters in batch and continuous mode with Corynebacterium glutamicum on lactic acid based on oxygen transfer and pH. Biotechnol Bioeng 98, 800–811. doi: 10.1002/bit.21359.

22. Grünberger A et al. 2013. Beyond growth rate 0.6: Corynebacterium glutamicum cultivated in highly diluted environments. Biotechnol Bioeng 110, 220–228. doi: 10.1002/bit.24616.

23. Wilks JC et al. 2009. Acid and base stress and transcriptomic responses in Bacillus subtilis. Appl Environ Microbiol 75, 981–990. doi: 10.1128/AEM.01652-08.

24. Slonczewski JL, Fujisawa M, Dopson M, Krulwich TA (Elsevier 2009), pp. 1–317.

25. Martinez KA et al. 2012. Cytoplasmic pH response to acid stress in individual cells of Escherichia coli and Bacillus subtilis observed by fluorescence ratio imaging microscopy. Appl Environ Microbiol 78, 3706–3714. doi: 10.1128/AEM.00354-12.

26. Zilberstein D, Agmon V, Schuldiner S, Padan E. 1984. Escherichia coli intracellular pH, membrane potential, and cell growth. J Bacteriol 158, 246–252. doi: 10.1128/JB.158.1.246-252.1984.

27. Liu Y et al. 2016. Mycothiol protects Corynebacterium glutamicum against acid stress via maintaining intracellular pH homeostasis, scavenging ROS, and S-mycothiolating MetE. J Gen Appl Microbiol 62, 144–153. doi: 10.2323/jgam.2016.02.001.

28. Peleg M, Normand MD, Corradini MG. 2005. Generating microbial survival curves during thermal processing in real time. J Appl Microbiol 98, 406–417. doi: 10.1111/j.1365-2672.2004.02487.x.

29. Mukamolova GV, Kaprelyants AS, Young DI, Young M, Kell DB. 1998. A bacterial cytokine. Proc Natl Acad Sci U S A 95, 8916–8921. doi: 10.1073/pnas.95.15.8916.

30. Hartmann M et al. 2004. The glycosylated cell surface protein Rpf2, containing a resuscitation-promoting factor motif, is involved in intercellular communication of Corynebacterium glutamicum. Arch Microbiol 182, 299–312. doi: 10.1007/s00203-004-0713-1.

31. Ryall B, Eydallin G, Ferenci T. 2012. Culture history and population heterogeneity as determinants of bacterial adaptation: the adaptomics of a single environmental transition. Microbiol Mol Biol Rev 76, 597–625. doi: 10.1128/MMBR.05028-11.

32. Bremer E, Krämer R. 2019. Responses of Microorganisms to Osmotic Stress. Annu. Rev. Microbiol. 73, 313–334. doi: 10.1146/annurev-micro-020518-115504.

33. Wood JM et al. 2001. Osmosensing and osmoregulatory compatible solute accumulation by bacteria. Comparative Biochemistry and Physiology Part A: Molecular & Integrative Physiology 130, 437–460. doi: 10.1016/S1095-6433(01)00442-1.

34. Novo D, Perlmutter NG, Hunt RH, Shapiro HM. 1999. Accurate flow cytometric membrane potential measurement in bacteria using diethyloxacarbocyanine and a ratiometric technique. Cytometry 35, 55–63. doi: 10.1002/(sici)1097-0320(19990101)35:1<55::aid-cyto8>3.0.co;2-2.

35. Miesenböck G, Angelis DA de, Rothman JE. 1998. Visualizing secretion and synaptic transmission with pH-sensitive green fluorescent proteins. Nature 394, 192–195. doi: 10.1038/28190.

36. Mahon MJ. 2011. pHluorin2: an enhanced, ratiometric, pH-sensitive green florescent protein. Adv Biosci Biotechnol 2, 132–137. doi: 10.4236/abb.2011.23021.

37. Wang P et al. 2010. Robust growth of Escherichia coli. Current biology 20, 1099–1103. doi: 10.1016/j.cub.2010.04.045.

38. Täuber S et al in preparation.

