## Supplement Information for "Growth response and recovery of *Corynebacterium glutamicum* colonies on single-cell level upon defined pH stress pulses"

**Table S1:** Single-cell sample size for each single stress pulse.

|  | **Stress pulse** | **Sample size** | **Growing cells** | **No growth** |
| --- | --- | --- | --- | --- |
| pH 4 | 30 min | 100 | 0 | 100 |
|  | 25 min | 119 | 5 | 114 |
|  | 20 min | 61 | 3 | 58 |
|  | 15 min | 50 | 6 | 44 |
|  | 10 min | 61 | 13 | 48 |
|  | 5 min | 45 | 41 | 4 |
| pH 5 | 2 h | 117 | 109 | 8 |
|  | 3 h | 94 | 54 | 35 |
|  | 4 h | 59 | 53 | 9 |
|  | 5 h | 117 | 37 | 80 |
|  | 6 h | 113 | 31 | 82 |
|  | 7 h | 128 | 11 | 117 |
|  | 8 h | 164 | 22 | 142 |
|  | 9 h | 152 | 9 | 143 |
| pH 10 | 2 h | 77 | 76 | 1 |
|  | 3 h | 82 | 67 | 15 |
|  | 4 h | 70 | 30 | 40 |
|  | 5 h | 108 | 32 | 76 |
|  | 6 h | 106 | 16 | 90 |
|  | 7 h | 72 | 12 | 52 |
| pH 11 | 5 min | 38 | 38 | 0 |
|  | 10 min | 53 | 44 | 9 |
|  | 15 min | 127 | 82 | 45 |
|  | 20 min | 181 | 146 | 35 |
|  | 25 min | 134 | 36 | 96 |
|  | 30 min | 95 | 0 | 95 |

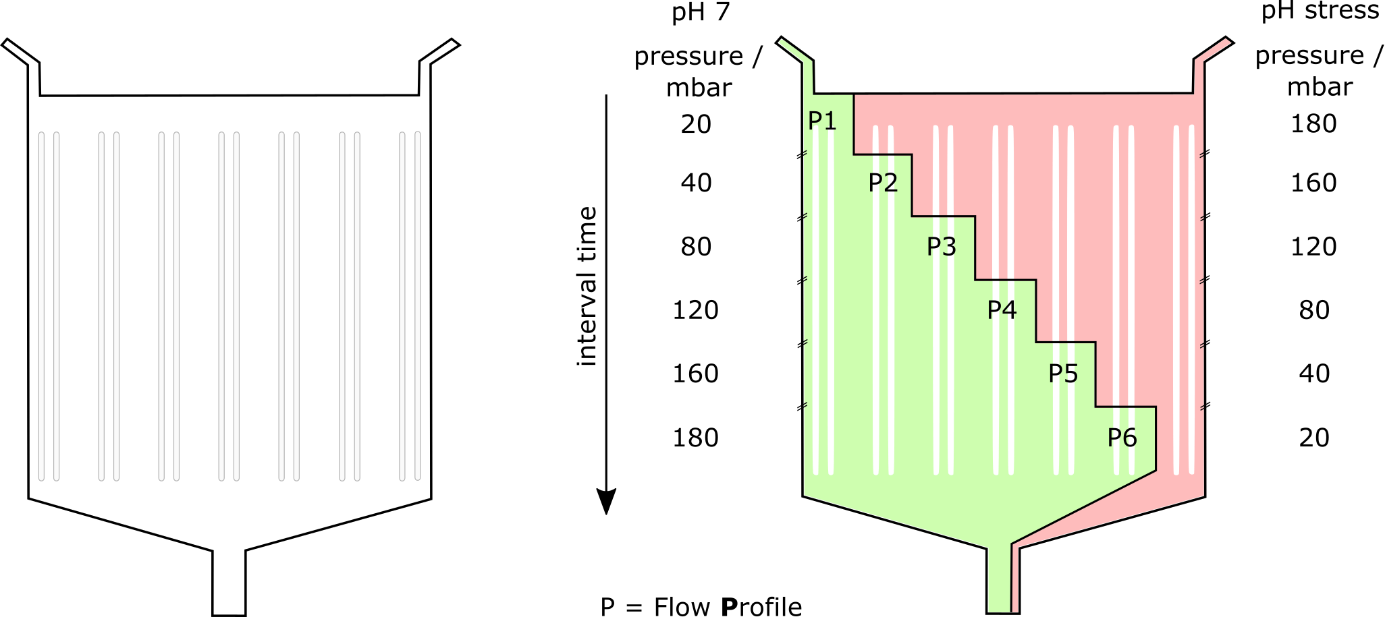

**Figure S1**: dMSCC with seven array pairs, the distance between these pairs being 400 µm. Six flow profiles were conducted with one control zone of each side.

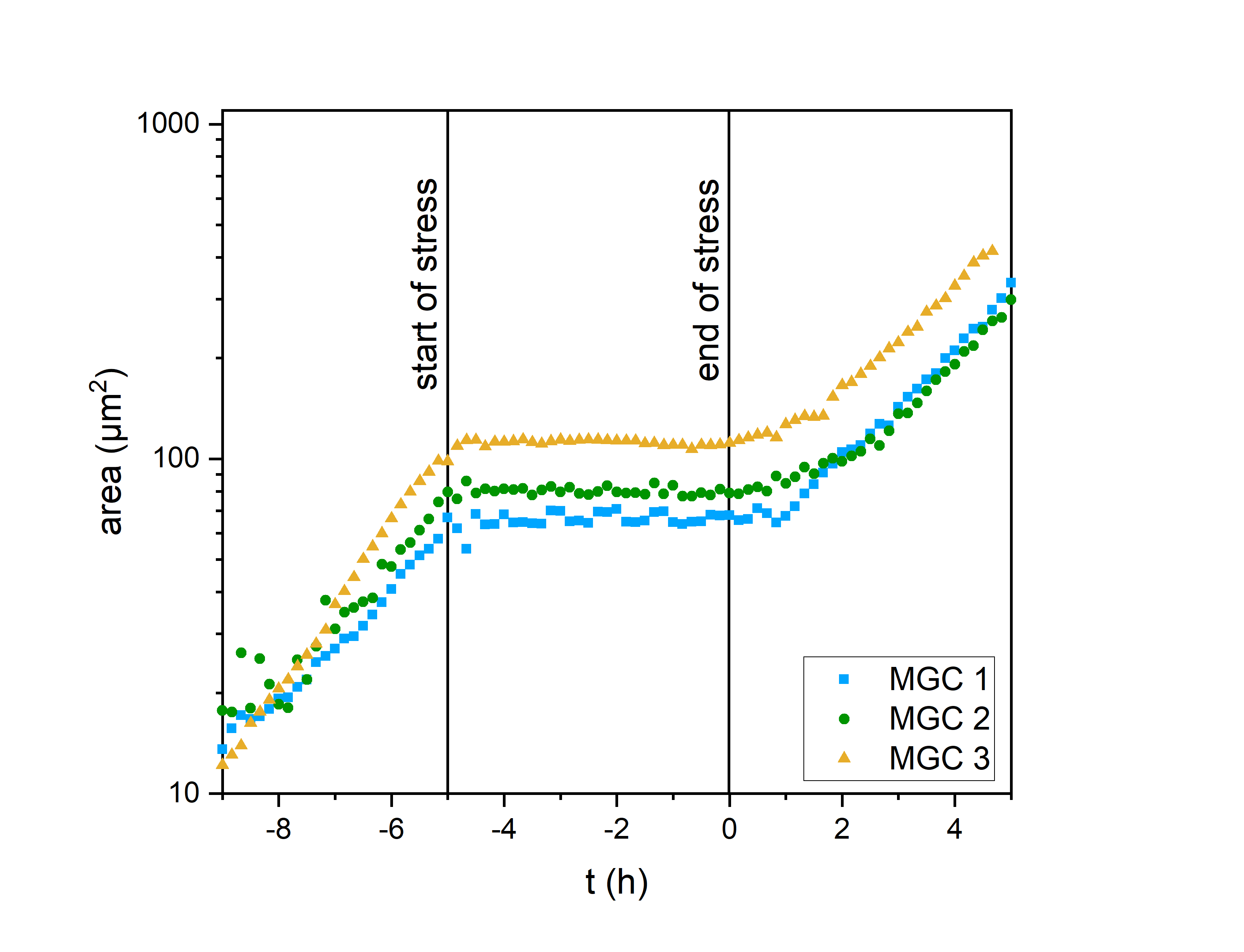

**Figure S2**: Growth curve at a single stress pulse of pH 5 for 5h. At t = 0h, the stress ends. Three colonies (red, green, blue) are shown.

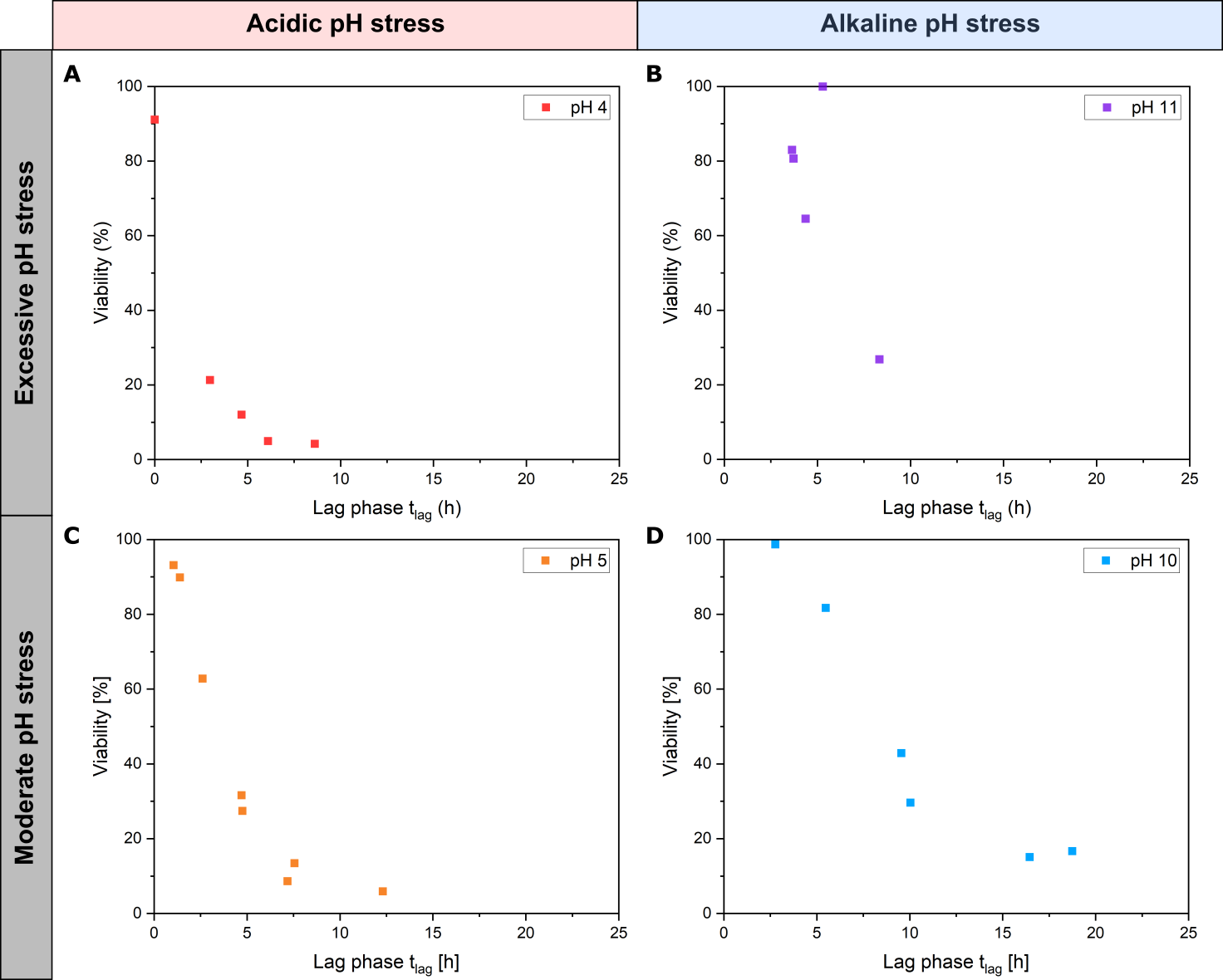

**Figure S3:** Correlation between viability and lag phase for colony growth of *C. glutamicum*. With decreasing viability, the lag phase increase for all pH values. A) pH 4. B) pH 11. C) pH 5. D) pH 10.

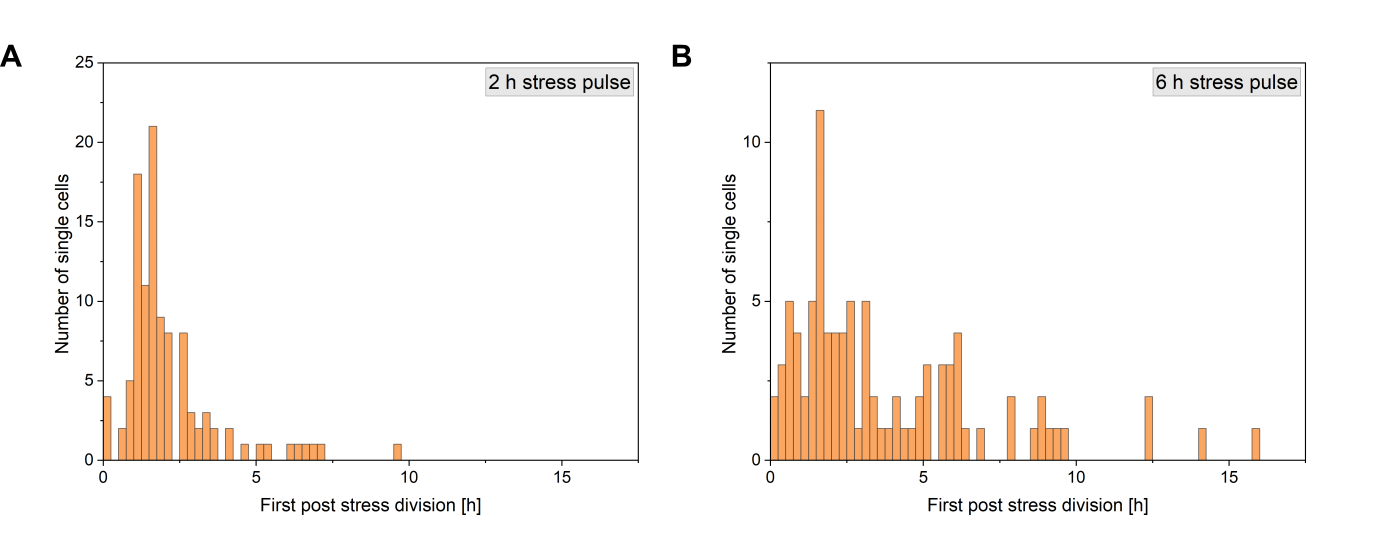

**Figure S4:** Cell distribution of the first post stress division time after a A) 2 h and B) 6 h pH 5 stress pulse.

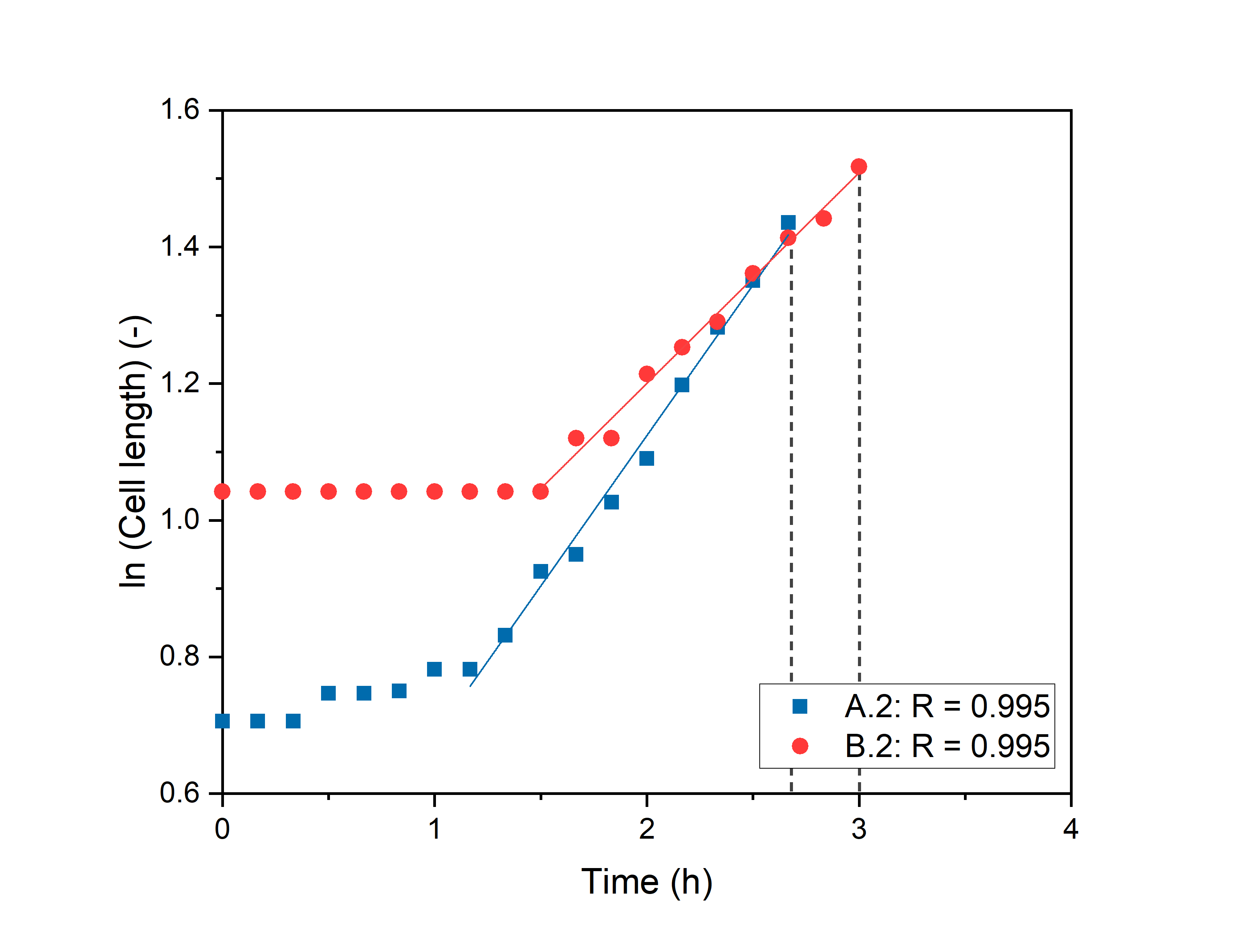

**Figure S5:** Correlation analysis for the exponential length growth of cell A.2 (2 h stress pulse at pH 5) and cell B.2 (2 h stress pulse of pH 10).

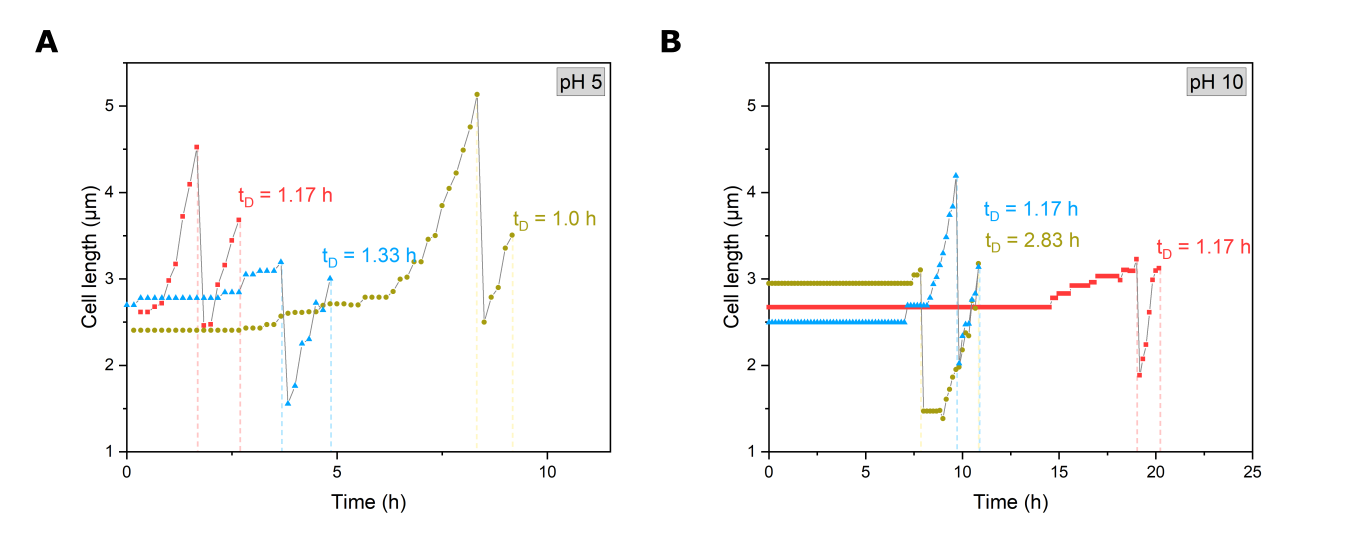

**Figure S6:** Cell length growth of single *C. glutamicum* cells after 6 h stress pulses of A) pH 5 and B) 10 stress until the first and second post-stress division. The dashed line marks the division of the cells.
